# Methods for analyzing continuous conformational variability of biomolecules in cryo electron subtomograms: HEMNMA-3D vs. traditional classification

**DOI:** 10.1101/2021.10.14.464366

**Authors:** Mohamad Harastani, Slavica Jonic

**Affiliations:** IMPMC - UMR 7590 CNRS, Sorbonne Université, MNHN 4 Place Jussieu, Paris, France

## Abstract

Cryogenic electron tomography (cryo-ET) allows studying biological macromolecular complexes in cells by three-dimensional (3D) data analysis. The complexes continuously change their shapes (conformations) to achieve biological functions. The shape heterogeneity in the samples imaged in the cryo electron microscope is a bottleneck for comprehending biological mechanisms and developing drugs. Low signal-to-noise ratio and spatial anisotropy (missing wedge artefacts) make cryo-ET data particularly challenging for resolving the shape variability. Other shape variability analysis techniques simplify the problem by considering discrete rather than continuous conformational changes of complexes. Recently, HEMNMA-3D was introduced for cryo-ET continuous shape variability analysis, based on elastic and rigid-body 3D registration between simulated shapes and cryo-ET data. The simulated motions are obtained by normal mode analysis of a high- or low-resolution 3D reference model of the complex under study. The rigid-body alignment is achieved via fast rotational matching with missing wedge compensation. HEMNMA-3D provides a visual insight into molecular dynamics by grouping and averaging subtomograms of similar shapes and by animating movies of registered motions. This article reviews the method and compares it with existing literature on a simulated dataset for nucleosome shape variability.

## 1. Introduction

Three-dimensional (3D) volumetric images of vitrified cell sections (slices) can be obtained using cryogenic electron tomography (cryo-ET). The 3D nature of cryo-ET data permits studying macromolecular complexes despite the crowded cell environment. The most common cryo-ET data acquisition scheme is the acquisition of multiple 2D projection images of a specimen rotated around a single axis (perpendicular to the electron beam) inside the electron mi-croscope. The obtained tilt-series 2D projections are used to reconstruct a 3D volume (referred to as tomogram) based on the Fourier slice theorem and backprojection. The tomogram of a cell section typically contains hundreds of copies of different macromolecules at random orientations. The copies of the macromolecule under study (target macro-molecule) can be identified and extracted into individual volumes called subtomograms, either manually or via template matching.

Subtomograms suffer from a low signal-to-noise ratio (SNR) due to exposing the sample to a low electron dose during data acquisition to preserve the fragile biological structure from radiation damage. Additionally, subtomograms suffer from spatial anisotropies, known as missing wedge artefacts, due to the inability to include in the 3D reconstruction the images from all orientations (the maximum tilt angle in the microscope is usually limited to ±60°), which corresponds to a missing wedge-shaped region in 3D Fourier space. The missing wedge artefacts are often observed as elongations along the beam axis, blurring and distracting caustics in the subtomograms. Due to the low SNR and the missing wedge artefacts, cryo-ET data processing is mainly based on rigid-body aligning and averaging of many subtomograms to enhance the data quality and reveal the target macromolecular structure, which is known as Subtomogram Averaging (StA) [1]. However, with recent instrumentation and software development, more research moves in the direction of studying subtomograms individually (e.g., development of methods for denoising and missing wedge correction with no or a minimum of averaging [2, 3]).

The main focus of studying macromolecules in their native state is to observe their shapes and dynamics. In the last decade, cryo-electron microscopy (cryo-EM) research has shown that disentangling macromolecular shape variability and identifying the macromolecular conformational transitions is valuable for understanding biological mechanisms [4, 5]. The most popular cryo-EM technique, namely Single Particle Analysis (SPA), allows a near-atomic resolu-tion of 3D reconstruction of macromolecules *in vitro* (from vitrified samples of a solution containing biochemically purified macromolecules), by combining alignment and classification in 2D and 3D [6]. Only some SPA methods consider continuous shape variability, meaning gradual conformational transitions of macromolecules with many intermediate conformational states. They represent images in a low-dimensional space allowing a 3D visualization of the macromolecular shape variability [7, 8]. In contrast to SPA, cryo-ET allows studying macromolecules *in situ* (in vitrified cell sections). The conformational variability in the native cellular environment is largely overlooked, due to the lack of cryo-ET image analysis methods capable of dealing with continuous shape variability in the data with low SNR and missing-wedge artefacts.

### Contribution

HEMNMA-3D [9] was introduced for cryo-ET continuous macromolecular shape variability analysis, inspired by HEMNMA, a method for continuous shape variability analysis in SPA [10, 8]. HEMNMA-3D is based on elastic and rigid-body 3D registration between simulated shapes and cryo-ET data. The simulated motions are obtained by normal mode analysis of a high- or low-resolution 3D reference model of the complex under study. The rigid-body alignment is achieved via fast rotational matching with missing wedge compensation. HEMNMA-3D provides a visual insight into molecular dynamics by grouping and averaging subtomograms of similar shapes and by animating movies of registered motions. This article reviews the method and compares it with existing literature on a simulated dataset for nucleosome shape variability.

## 2. Related work

### 2.1. 3D elastic registration for medical images

Elastic registration is extensively used in medical image processing for a wide spectrum of applications, including Magnetic Resonance Imaging [11] and Computed Tomography [12]. Noteworthy, several multi-purpose machine learning and deep learning-based 3D registration methods were proposed [13, 14]. However, none of these methods was adapted or applied to cryo-ET, possibly due to severe cryo-ET data limitations such as missing wedge artefacts and poor SNR.

### 2.2. Cryo-ET subtomogram classification

Methods reported to deal with cryo-ET data heterogeneity are based on classification and rigid-body alignment. They can be split into two categories, namely multireference alignment approaches [15] and post-alignment classification approaches [16].

Multi-reference alignment approaches are based on competitive alignment. An expert user provides a number of ref-erences with different shapes of the macromolecule under study based on anticipated shape variabilities. An algorithm then aligns (rigid-body-wise) and compares each subtomogram with the multiple given references and attributes it to the reference that yields the highest similarity score. The starting references evolve by averaging the aligned subtomograms and repeating the process until stability. Such methods require prior knowledge of the biological specimen and are prone to overfitting and data misinterpretation if not carefully used.

Post-alignment classification approaches perform classification after rigid-body alignment via StA. During StA, subtomograms are rigid-body aligned against a reference to maximize a scoring function. In each iteration, the aligned subtomograms are averaged to produce a structure that becomes the reference for the next StA iteration. The iterations repeat until convergence. The result is an average structure and aligned subtomograms (centred and oriented with respect to the reference used in the last iteration). Classifying aligned subtomograms according to the shape variability is challenging because of the missing wedge artefacts. The aligned subtomograms are usually classified based on the covariance matrix calculated using a constrained correlation coefficient (CCC) between each pair of aligned subtomograms. The CCC corresponds to constraining the cross-correlation evaluation to the Fourier-space region that excludes the missing wedge of both subtomograms [17]. The covariance matrix serves as a basis for a hierarchical classification technique, or it is fed to a dimensionality reduction method first and, then, to a clustering technique [17, 18]. A drawback of post-alignment classification methods is their dependency on rigid-body alignment accuracy, which decreases as the specimen’s conformational heterogeneity increases.

In practice, the classification remains efficient at resolving discrete structural variabilities such as ligand binding and macromolecular disassembly, but for classifying continuous shape variants, particles assigned to the same class will rarely, if ever, have identical conformations.

## 3. HEMNMA-3D

A flowchart for HEMNMA-3D is in Figure 1. The method comprises the following blocks:

1. Input: subtomograms and a **reference structure**;
2. **Normal Mode Analysis** of the reference structure;
3. **Elastic and rigid-body 3D registration**; and
4. **Visualizing macromolecular shape variability** via selective data averaging and animations.

**Figure 1.**
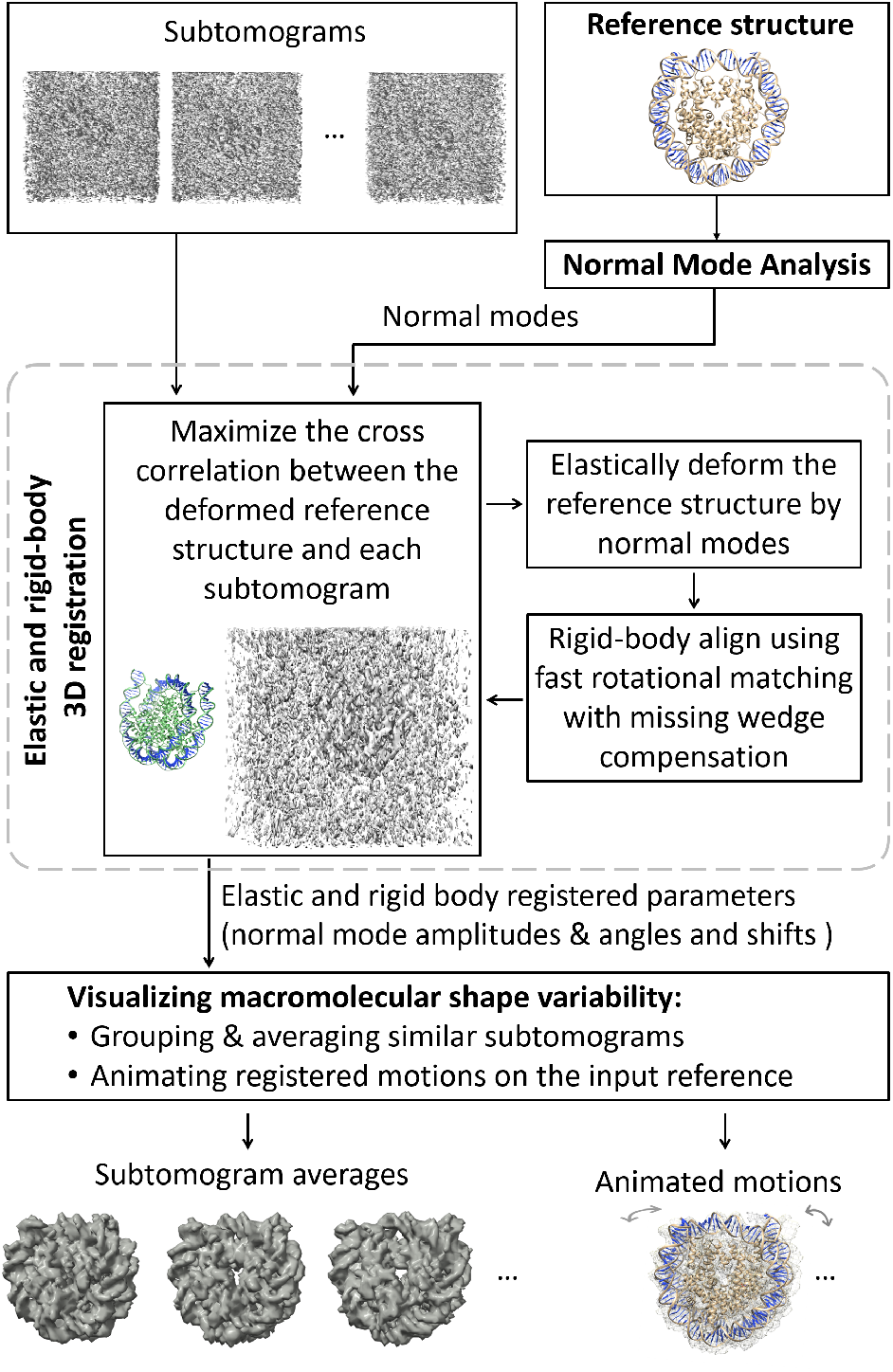
Flowchart of HEMNMA-3D for macromolecular shape variability analysis in cryo-ET using 3D elastic registration via normal mode analysis. Note: the displayed molecule is the nucleosome (PDB:3w98).

### 3.1. Reference structure

The 3D registration is performed using a reference structure in the Protein Data Bank (PDB) format. If there is an existing atomic model of the macromolecule under study (e.g., a model available in the PDB database and obtained by X-ray crystallography, nuclear magnetic resonance, or derived from 3D reconstructed density maps from cryo-EM images), this model can be used as the reference structure for the method. However, if no atomic model is available but a lower-resolution density map can be obtained by SPA reconstruction or subtomogram averaging, this density map can be converted into a pseudoatomic model [19, 20] and, then, this model can be used as the reference structure for the method. The PDB file of the reference structure contains 3D Cartesian coordinates of atoms or pseudoatoms.

### 3.2. Normal Mode Analysis

Normal Mode Analysis (NMA) is a method for molecular mechanics simulation. One of its main applications is the elastic deformation of an existing atomic structure to fit a cryo-EM density map of the same macromolecule but at a different conformation. This application is usually known as normal mode flexible fitting and allows deriving atomic models from cryo-EM maps [21].

Normal modes are computed from the atomic or pseudoatomic reference structure based on the elastic network model [22]. This model represents the atoms or pseudoatoms as locally connected (within a cutoff distance) by elastic springs. Normal modes are vectors that describe harmonic-oscillator motions of the elastic network model (the length of each normal-mode vector is 3 times the number of atoms or pseudoatoms). Computationally, normal modes are eigenvectors of a Hessian matrix of the system (the matrix of second derivatives of the potential energy function of the elastic network model) and the corresponding eigenvalues are the squares of the normal-mode frequencies. Also, for each normal mode, its collectivity degree is computed, which measures the percentage of atoms or pseudoatoms that move with that mode. Low-frequency, high-collectivity normal modes have been shown to be relevant to macromolecular conformational variability [23].

The atomic or pseudoatomic displacement is determined by a linear combination of normal modes, where the coefficients of the linear combination are the displacement amplitudes along the modes. NMA only allows computing normal modes (vectors), meaning atomic or pseudoatomic displacement directions. The amplitudes of these displacements are computed by 3D elastic registration. In the context of 3D elastic registration, the key advantage of NMA is that selecting a subset of normal modes with the lowest frequencies and highest collectivities allows faster data analysis and a regularization against noise overfitting, which was also observed in 3D-to-2D elastic registration [8].

### 3.3. Elastic and rigid-body 3D registration

This module comprises simultaneous NMA-based elastic registration and Fast Rotational Matching (FRM)-based rigid-body alignment of the reference structure with each given subtomogram via numerical optimization.

Given a subtomogram, a reference structure, and a set of normal modes of the reference structure, this module searches for the displacement amplitudes along the normal modes (elastic parameters) and the angles and shifts (rigid-body parameters) of the reference structure to match the structure to the subtomogram. A numerical optimizer maximizes the similarity between the subtomogram and a density volume simulated [24] from the elastically deformed, oriented, and shifted atomic or pseudoatomic reference. The similarity measure is the CCC, which is here defined as the cross-correlation between the reference and subtomogram density maps excluding the Fourier-space region corresponding to the subtomogram’s missing wedge. Such similarity measure allows to compensate for the missing wedge, which otherwise can lead to erroneous results. The numerical optimizer is a variant of Powell’s UOBYQA method known as CONDOR [25], which subjects the objective function to a trust-region radius and avoids noise over-fitting by giving more credibility to smaller normal mode amplitudes during the registration.

For each subtomogram, the corresponding normal mode amplitudes are initiated with zeros, i.e. the non-deformed reference is used in the first iteration. As the iterations evolve, the reference model is deformed with the new guesses of the normal mode amplitudes, converted into a volume and rigid-body aligned with the subtomogram using FRM [26]. The rotational matching allows a fast and accurate 6D search for orientations and shits (three Euler angles and three translation parameters). At the end of each iteration, the CCC is found and fed to the numerical optimizer. The iterations repeat, refining the elastic and rigidbody registration parameters until convergence.

### 3.4. Macromolecular shape variability visualization

The number of elastic registration parameters (normal-mode amplitudes) is determined by the number of normal modes used for the elastic registration. If more than three normal modes were used, the ensemble of normal-mode amplitudes (for all subtomograms) can be projected onto a lower-dimensional space using a dimensionality reduction technique, e.g. Principal Component Analysis (PCA). The obtained high- or low-dimensional space of normal mode amplitudes (conformational space) allows a global data display for interpreting macromolecular shape variability. Each point in this space represents a subtomogram, and close points correspond to similar registered shapes. In this conformational space, the shape variability can be analyzed in the following two ways: 1) by averaging subtomograms of similar registered shapes; and 2) by animating the registered motion of the reference structure along different directions.

#### Grouping and averaging similar subtomograms

As close points in the conformational space correspond to similar registered shapes, grouping and averaging close points in dense regions can help visualize variable macromolecular shapes with better SNR and less artefacts (attenuated noise and missing wedge artefacts thanks to averaging of many similar shapes). Before computing the group averages, the rigid-body alignment parameters found via the elastic and rigid-body 3D registration module are applied to the subtomograms. The subtomogram averages from different regions of the conformational space can be overlapped for a visual comparison.

#### Animating registered motions of the input reference

We can further analyze the conformational space by animating registered motions for several points through data distribution manifolds (e.g., following a straight line or a curve fitting the data). If a dimensionality reduction technique is applied, the inverse mapping must be first used (e.g. inverse PCA) to find the corresponding normal mode amplitudes for each point. These normal modes amplitudes are applied to elastically deform the reference structure (one elastically deformed structure is obtained for each selected point). By concatenating the resulting structures, a movie-like animation of registered motion can be obtained.

## 4. Implementation

The software of HEMNMA-3D is open-source and available on GitHub [10, 9]. It is built as part of the Continuous-Flex plugin of the open-source Scipion software package [27], commonly used for cryo-EM data processing. Our software provides a graphical user interface (GUI) and is empowered with a C++ backend with a message passing interface (MPI) parallelization scheme. The number of subtomograms that can be processed simultaneously depends on the number of available CPUs. The current implementation takes around 10 minutes to analyze a subtomogram of size 643 voxels using three normal modes (tested on 2.2 GHz Intel Xeon Silver 4214 CPU processor and 64 GB RAM). The more normal modes are used, the slower the processing.

## 5. Experiment

### 5.1. Simulating nucleosome shape variability

To review and compare HEMNMA-3D performance with existing literature, we synthesized a dataset comprising 1000 subtomograms with an imagined continuous shape variability of the nucleosome. We generated a linear combination of two reported motions for the nucleosome, breathing and gaping [28], with a linear dependence between the amplitudes of normal modes corresponding to the two motions.

First, we performed NMA of the nucleosome atomic structure available in the PDB database under the code 3w98 and we visualized the motions carried by the different computed normal modes. Among these modes, we identified the modes describing breathing and gaping motions as normal modes 9 and 13, respectively (note here that the mode number corresponds to the frequency of the mode and that higher numbers correspond to higher frequencies). We generated a dataset using a linear relationship between the amplitudes of normal modes 9 and 13 so that the nucleosome is simultaneously breathing and gaping. Precisely, at one end of the generated ground-truth conformational distribution, the nucleosome’s two DNA ends (arms) are moving away from each other, and at the same time, the gap between the two DNA gyres increases. At the other end of the generated conformational distribution, the DNA arms approach each other and the gap between the two DNA gyres decreases. We simulated a gradual transition between the two ends, representing a continuum of nucleosome shapes, combining breathing and gaping. Equal random amplitudes uniformly distributed in the range [−150, 150] were used for the two normal modes 9 and 13. An illustration of the simulated movements is provided in Figure 2.

**Figure 2.**
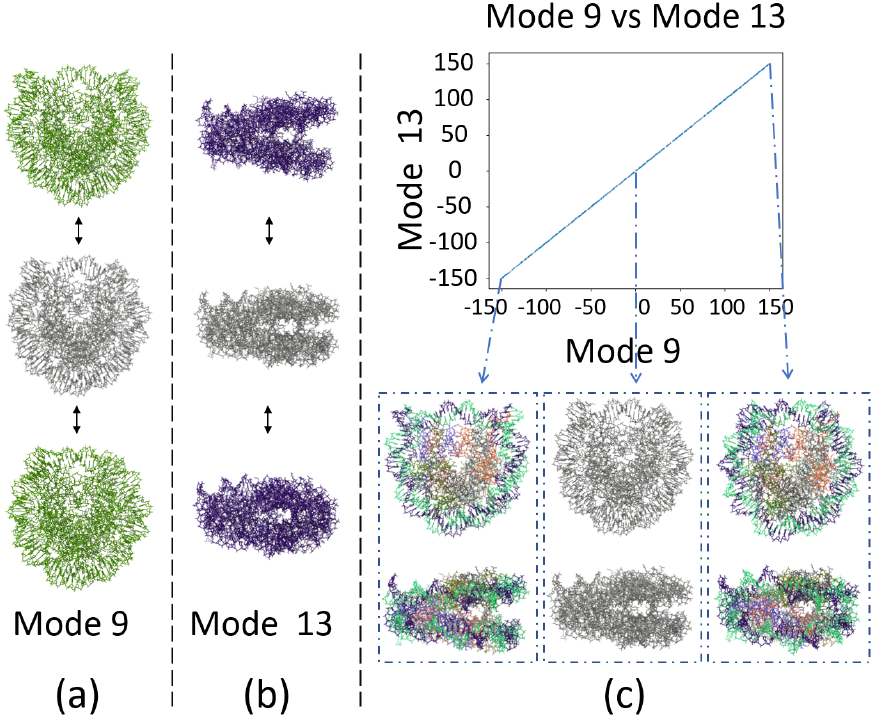
Synthesized combined breathing and gaping motions of the nucleosome (PDB 3w98 structure): (a) nucleosome breathing motion, (b) nucleosome gaping motion, (c) generated groundtruth conformational distribution (top) comprising 1000 synthetic nucleosome shape variants obtained by a linear combination of modes 9 and 13, with a linear dependence between the normal-mode amplitudes (blue points in the plot), and 3 representative shapes (bottom) corresponding to the two ends and the middle of the conformational distribution.

To generate this dataset, for each subtomogram, we performed the following steps:

1. Elastically deform the atomic structure (PDB:3w98) using equal random amplitudes for the two normal modes 9 and 13 in the range [−150, 150].
2. Convert the elastically deformed structure to a density map of size 64 × 64 × 64 voxels (voxel size: 3.45 Å × 3.45 Å × 3.45 Å), using [24].
3. Rotate and shift the volume in 3D space using random Euler angles and random x, y, z shifts (the random shift range is ±5 pixels from the center).
4. Tilt and project the randomly rotated and shifted volume, using the tilt angle from −60° to +60° with 1° step, to obtain a collection of 2D projection images (i.e. tilt series).
5. Simulate microscope conditions by adding heavy noise (SNR = 0.01) and modulating the images with the contrast transfer function (CTF) of the microscope (using the defocus of −0.5 μm).
6. Invert the CTF phase (a common CTF correction).
7. Reconstruct a volume (our synthetic subtomogram) from the obtained tilt series using a Fourier reconstruction method [29].

Figure 3 shows an example subtomogram from the synthesized dataset and the corresponding ideal volume, for comparison in real space and in Fourier space.

**Figure 3.**
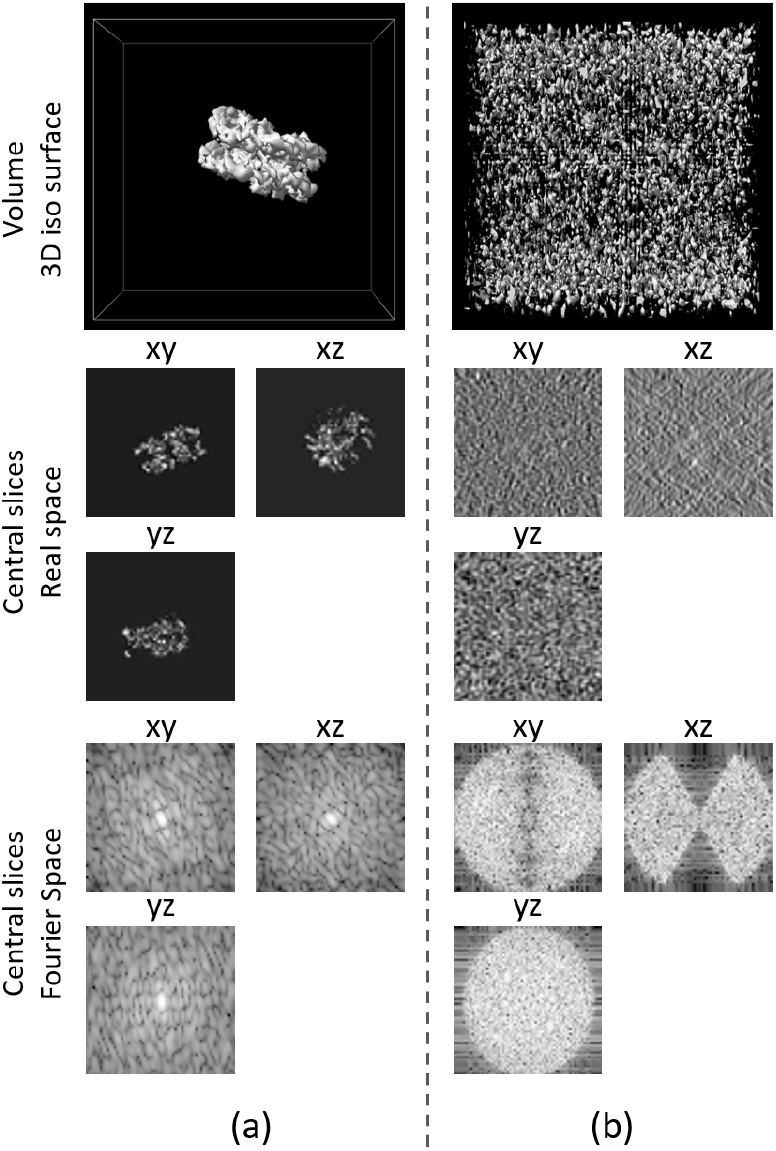
Example of a noisy and missing-wedge affected synthetic subtomogram compared with the corresponding ideal volume of the nucleosome: (a) ideal volume (without noise and without missing wedge artefacts), (b) noisy and missing-wedge affected synthetic subtomogram.

### 5.2. Traditional subtomogram averaging and post alignment classification

StA provides a global average without considering the shape variability, and it provides a basis for performing classification of subtomograms (classification of the subtomograms aligned through StA).

We applied StA on the synthesized nucleosome dataset, using the protocol based on the rigid-body alignment approach of [26] (recall that this rigid-body alignment approach is also used in the elastic and rigid-body alignment of HEMNMA-3D). This StA protocol uses an exhaustive angular search (with FRM method) and a shifts search within a region of interest, and compensates for the missing wedge by using the CCC (evaluation of the correlation between the subtomogram average of each iteration and the given subtomogram density maps, but excluding the evaluation in the missing-wedge region of the given subtomogram).

We followed the procedure in [26] and set the shifts search region to 10 voxels from the image center. We started iterations using an average of the unaligned subtomograms (this StA procedure is referred to as reference-free alignment). After six iterations, StA converged (further iterations gave the same results). The StA averages are shown in Figure 4.

**Figure 4.**
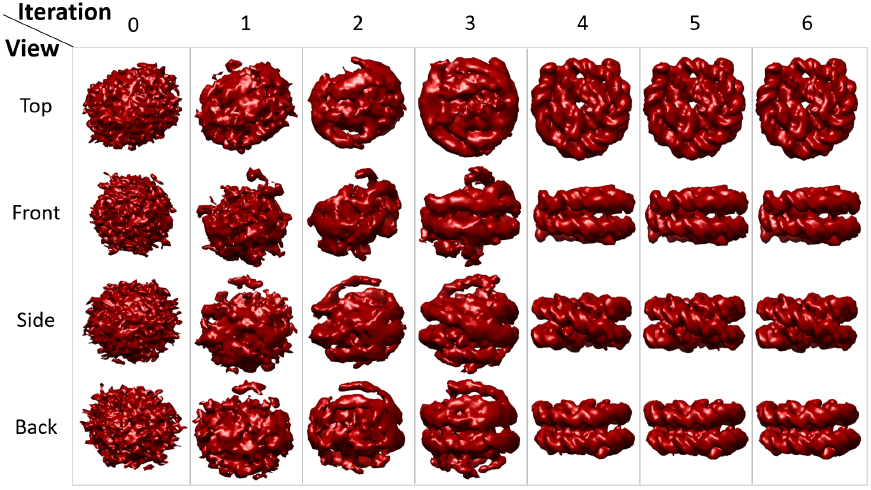
Subtomogram averaging applied to the synthesized nucleosome subtomograms. A reference-free alignment was performed using Fast Rotational Matching [26].

After StA, we applied the obtained rigid-body alignment parameters (found through StA) on the subtomograms, and we evaluated the covariance matrix CCC_ij_ of pairwise constrained cross-correlation (see section 2.2 for more details). We performed the two most common post-alignment classification techniques on the CCC_ij_ matrix, namely hierarchical clustering [18] and PCA followed by k-means [17].

The hierarchical clustering on 1-CCC_ij_ matrix was performed to 10 classes using the Agglomerative Clustering module of Python Scikit-Learn package (version 0.22.1 and default parameters were used) [30]. We note that applying the clustering algorithm directly on the CCC_ij_ matrix gives identical results, and we used the convention proposed in the literature [18, 17]. The clustering tree (dendrogram) and class averages are shown in Figure 5.

**Figure 5.**
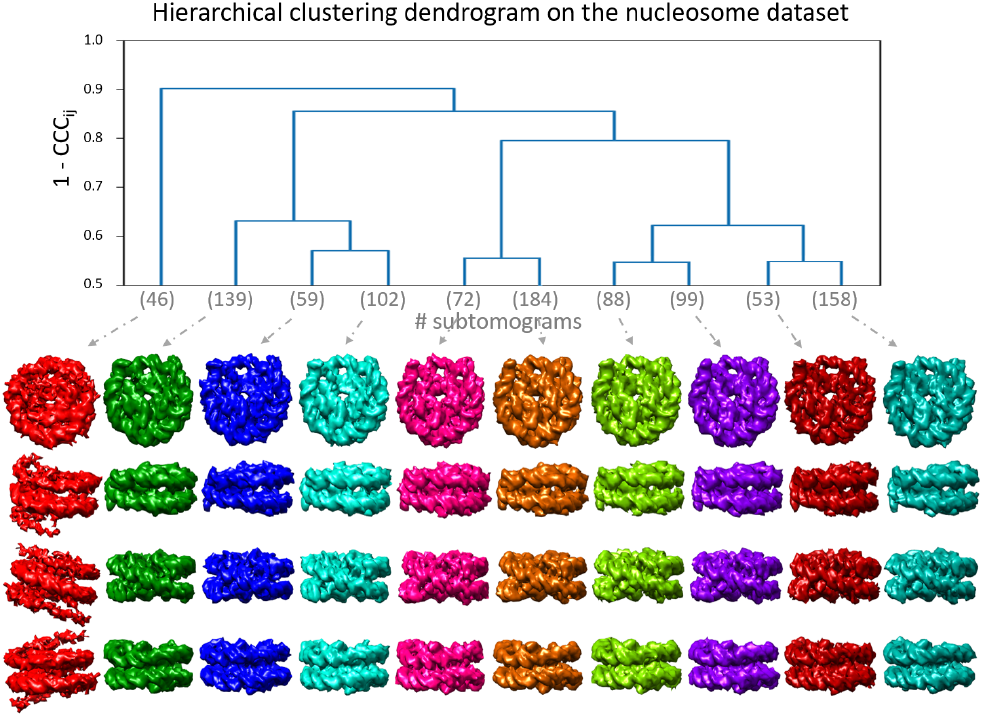
Hierarchical clustering applied to the synthesized nucleosome subtomograms. Top: hierarchical tree for 1-CCC_ij_ matrix. Bottom: views (vertically in the same color) of different subtomogram class averages (horizontally in different colors).

The k-means clustering was performed following PCA on the CCC_ij_ matrix. The clustering was done into 10 classes (k=10) based on the first two principal axes, using the k-means module of Scikit-Learn. In general, the choice of the number of principal axes to perform classification is arbitrary, as explained in [17]. Since the dataset was synthesized with two degrees of freedom (nucleosome breathing and gaping), we set the number of principle axes to 2, to obtain the best results. Figure 6 shows the classification of the PCA space and the resultant class averages.

**Figure 6.**
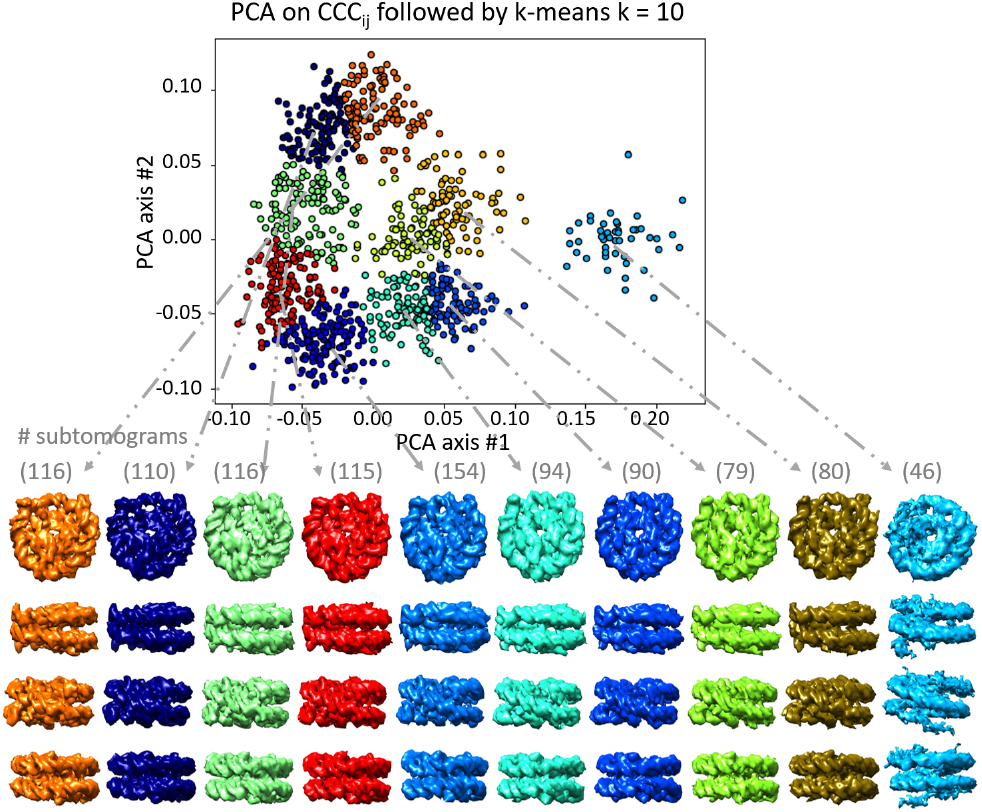
K-means clustering applied to the synthesized nucleosome subtomograms. Top: k-means clustering in the space of the first two PCA axes of CCC_ij_ matrix. Bottom: views (vertically in the same color corresponding to the color in the PCA space) of different subtomogram averages (horizontally in different colors).

We note that the two tested classification techniques give similar outputs, showing different discrete class averages of the nucleosome, at different breathing and gaping magnitudes (Figures 5 and 6). However, these outputs do not allow an unambiguous interpretation of the results in terms of the synthesized ground-truth conformational transitions of the nucleosome (from the smallest magnitudes to the largest magnitudes of breathing and gaping and vice versa).

### 5.3. HEMNMA-3D

Applying HEMNMA-3D to the synthesized nucleosome dataset aims at solving the inverse problem of finding the nucleosome shape variant in each subtomogram, i.e. estimating the amplitudes of normal modes 9 and 13 of the PDB structure 3w98 as close as possible to the generated ground-truth amplitudes. We set the method parameters as follows:

- **NMA settings:** To make the elastic and rigid-body 3D registration task more realistic and challenging, we used three normal modes (modes 9, 10 and 13) instead of only two modes (modes 9 and 13 used to generate the dataset).
- **FRM settings** The shift range for the rigid-body registration (FRM method) is set to 10 pixels.

The amplitudes estimated for modes 9 and 13 using HEMNMA-3D are shown in Figure 7 (a). It is graphically intelligible that the linear relationship is retrieved between the estimated amplitudes of the two modes. Figure 7 (b) shows the histogram of the amplitudes estimated for mode 10 and confirms that they are globally near zero.

**Figure 7.**
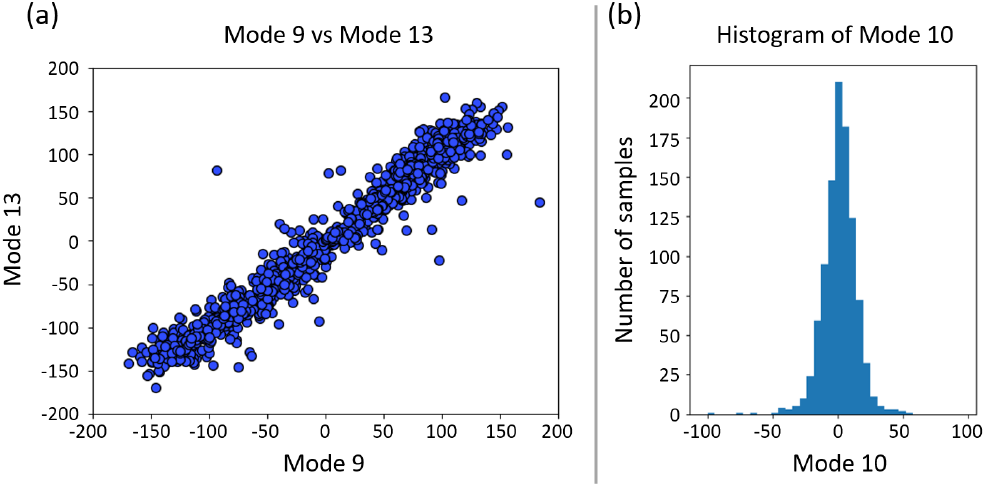
Output of the elastic and rigid-body 3D registration module of HEMNMA-3D using synthesized nucleosome subtomograms. The goal was the retrieval of the ground-truth amplitudes of normal modes 9, 10 and 13. Ideally, the amplitudes of mode 10 are equal to zero and there is a linear relationship between the amplitudes of modes 9 and 13 in the range [−150, 150]: (a) amplitudes of mode 9 vs amplitudes of mode 13, (b) histogram of amplitudes of mode 10.

Table 1 presents the mean absolute error between the estimated and ground-truth normal-mode amplitudes and the standard deviation of the error. It should be noted that 14/1000 points were excluded from the statistics as found to differ significantly (outlier points) from the remaining observations. These points were excluded for having a p-value below 10^−4^ based on the Mahalanobis distance [31].

**Table 1.**
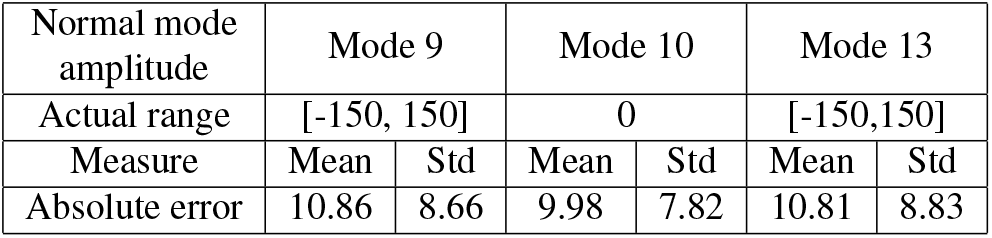
Mean absolute error between the estimated and groundtruth normal-mode amplitudes and the standard deviation of the error, for HEMNMA-3D using synthesized nucleosome subtomograms. Points below the p-value of 10^−4^ were excluded (14/1000 points) from the error evaluation based on the Mahalanobis distance.

Normal-mode amplitudes do not have a physical unit. Nonetheless, the Root Mean Square Deviation (RMSD) [32] between the reference atomic coordinates and these coordinates displaced using the calculated errors as the normal-mode amplitudes can transform these errors in physical units. The nucleosome core complex comprises eight histone proteins surrounded by 146 DNA base pairs. The synthesized movements (breathing and gaping) mainly impacted the DNA loops. Evaluating the RMSD without excluding the core histones can give a false sense of achieving higher accuracy by pulling the RMSD value towards zero. Therefore, the reported RMSD hereafter is based on the nucleosome’s DNA loops only (chain I and J of the PDB structure 3w98).

We found a RMSD of 0.44 Å corresponding to the mean absolute errors in Table 1 (for a combined displacement along modes 9, 10 and 13). Also, we found a RMSD of 0.79 Å corresponding to the sum of the mean and standard deviation of the errors in Table 1. Hence, the error range is significantly inferior to the pixel size used to create the data (3.45 Å).

Figure 8 (a) shows grouping and averaging of subtomograms through the point distribution in the conformational space (ten equally distanced groups). The corresponding subtomogram averages show the expected combination of continuous motions of breathing and gaping, which can be compared with the ground-truth motion in Figure 2.

**Figure 8.**
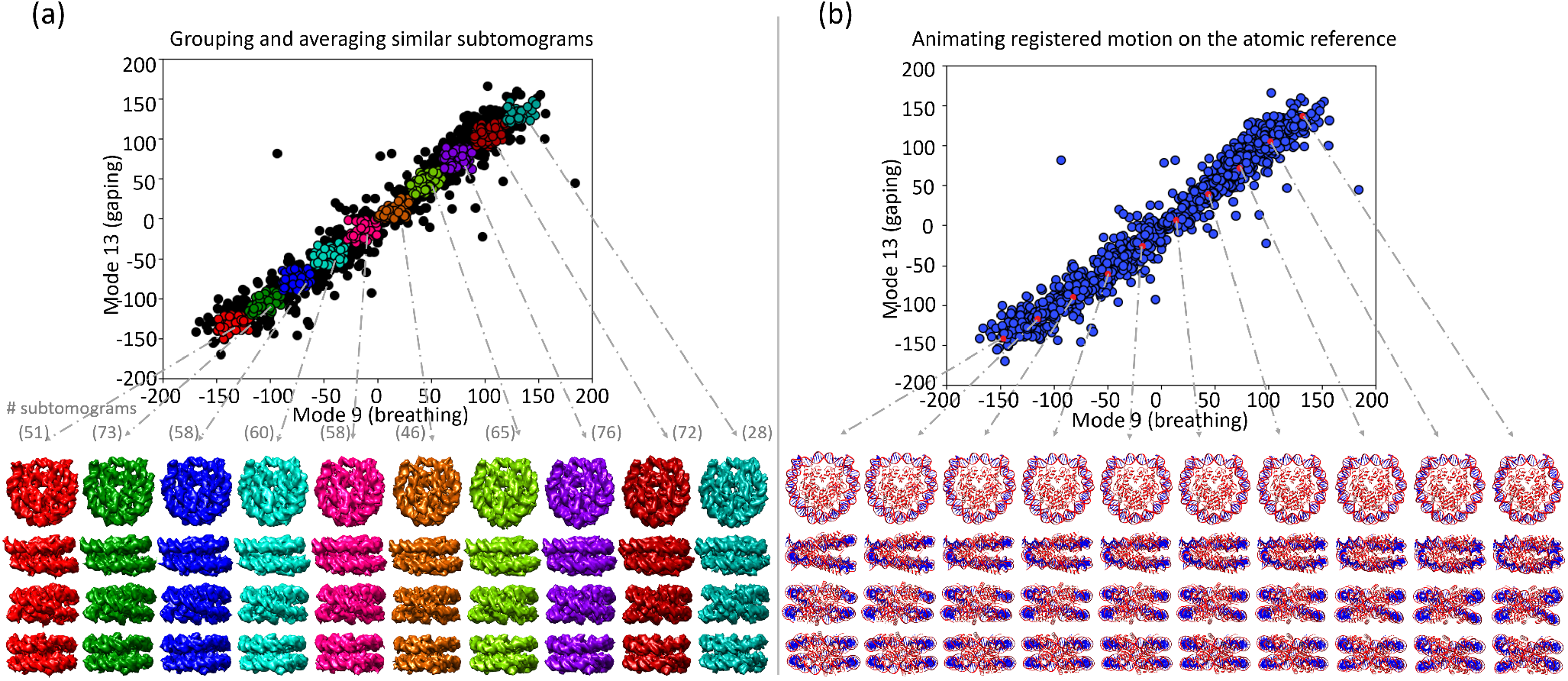
HEMNMA-3D applied to the synthesized nucleosome subtomograms. (a) group averages for ten equally distanced groups along the subtomogram (point) distribution in the conformational space, (b) displacement of the reference PDB structure 3w98 along the direction of the data distribution in the conformational space, 10 frames represented by red dots. Note: each column represents four different views of the same structure.

Figure 8 (b) shows the displacement of the reference structure along 10 points in the direction of the point distribution in the conformational space (the procedure explained in Section 3.4).

The obtained subtomogram averages and animation show that the ground-truth nucleosome motion (a combination of breathing and gaping) was retrieved.

## 6. Discussion and conclusion

This article presented a review on HEMNMA-3D, a method that addresses continuous macromolecular shape variability captured in cryo-ET subtomograms. HEMNMA-3D applies elastic and rigid body 3D registration via normal mode analysis, fast rotational matching and trust-region based numerical optimization.

We tested HEMNMA-3D by synthesizing a dataset of nucleosome shape variability under challenging conditions of the simulated microscope and testing the method’s capability to recover the ground-truth shapes. The test results indicate that the method recovers the ground-truth combination of two nucleosome motions (breathing and gaping) with subpixel accuracy. Also, it does not overfit the data with non-existing motions (normal mode 10 was not used to create the nucleosome dataset but used for 3D elastic registration, and it was retrieved with amplitudes near zero).

We applied two state-of-the-art methods for cryo-ET classification on the synthesized nucleosome dataset. Both methods gave similar outputs, showing different discrete class averages of the nucleosome at different breathing and gaping magnitudes. However, the choice of the number of classes is arbitrary in these methods and the shape transitions between the obtained class averages are ambiguous, probably because of the continuous nature of the shape variability.

In contrast to these methods, HEMNMA-3D adopts a new scheme that permits revealing hidden macromolecular dynamics by i) grouping and averaging similar subtomograms at locations in the conformational space that reveal the shape transitions and ii) animating the reference structure by displacing it in different directions in the conformational space. Hence, HEMNMA-3D provides a promising new insight into what can be achieved in cryo-ET studies of macromolecular shape variability. For more details on HEMNMA-3D and an example of its use and results with experimental cryo-ET subtomograms, the reader is referred to [9].

However, unlike the classification methods, HEMNMA-3D is limited to macromolecular elastic shape variability that can be explained with NMA. It is not suitable for analyzing other structural variabilities such as macromolecular disassembly or binding and unbinding of ligands. Future work can involve combining this method with classification to first disentangle such discrete structural variabilities and then analyze continuous intraclass variability.

## 7. Acknowledgements

We acknowledge the support of the French National Research Agency - ANR (ANR-20-CE11-0020-03 and ANR-19-CE11-0008-01 to S.J.); the Sorbonne University (2019 “Interface pour le Vivant” PhD scholarship grant to M.H.); and the access to the HPC resources of CINES and IDRIS granted by GENCI (A0100710998, A0070710998, AP010712190, AD011012188 to S.J.).

